# Spatial computing for the control of working memory

**DOI:** 10.1101/2020.12.30.424833

**Authors:** Mikael Lundqvist, Scott L Brincat, Jonas Rose, Melissa R. Warden, Tim Buschman, Earl K. Miller, Pawel Herman

**Author notes:** co-last authors (Earl K. Miller, Pawel Herman).

## Abstract

Working memory (WM) allows us to selectively remember and flexibly control a limited amount of information. Earlier work has suggested WM control is achieved by interactions between bursts of beta and gamma oscillations. The emerging question is how beta and gamma bursting, reflecting coherent activity of hundreds of thousands of neurons, can underlie selective control of individual items held in WM? Here, we propose a principle for how such selective control might be achieved on the neural network level. It relies on spatial computing, which suggests that beta and gamma interactions cause item-specific activity to flow spatially across the network over the course of a task. This way, control-related information about, for instance, item order can be retrieved from the spatial activity independent of the detailed recurrent connectivity that gives rise to the item-specific activity itself. The spatial flow should in turn be reflected in low-dimensional activity shared by many neurons. We test predictions of the proposed spatial computing paradigm by analysing control-related as well as item-specific activity in local field potentials and neuronal spiking from prefrontal cortex of rhesus macaques performing four WM tasks. As predicted, we find that the low-dimensional activity has a spatial component from which we can read out control-related information. These spatial components were stable over multiple sessions and did not depend on the specific WM items being used. We hypothesize that spatial computing can facilitate generalization and zero-shot learning by utilizing spatial component as an additional information encoding dimension. This offers a new perspective on the functional role of low-dimensional activity that tends to dominate cortical activity.

## Introduction

Working memory (WM) is a mental sketchpad for the short-term storage and top-down control of information (Baddeley, 1992; Miller & Cohen, 2001, D’esposito et al., 1995; 2000; Goldman-Rakic, 1996; DeNicola et al., 2020; Oberauer, 2002; Sauseng et al., 2005). This control is key to WMs central role in cognition (Goldman-Rakic, 1996). We can select what we retain, read out or delete from WM as well as manipulate the contents (Oberauer, 2002; Sauseng et al., 2005; Warden and Miller, 2010; Chatham et al., 2014; Wolff et al., 2017; Yu et al., 2020; Lewis-Peacock et al., 2015; Lundqvist et al., 2018; van Ede et al., 2017; Miller et al., 2020). It is not clear however what mechanisms underlie such flexible control.

Recent work on the temporal dynamics of WM have begun to provide some insight. The central idea is that top-down control stems from interactions between bursts of gamma and beta power (Lundqvist et al., 2011; 2016; 2018; Miller et al., 2018). The gamma bursts are associated with spiking that in turn encodes and maintains WM content. Beta bursts act as the control signal. They carry top-down information and inhibit gamma/spiking, thus controlling the access to WM contents carried by the spiking. This is supported by empirical observations of an anti-correlated “push-pull” relationship between beta and gamma during the encoding, read-out, and deletion of the contents of WM (Lundqvist et al., 2018). For example, when information is encoded into, or read out from, WM, beta decreases and gamma increases along with spiking carrying the WM content. When the content is no longer relevant and could be cleared out of the WM, the opposite dynamics is seen (Lundqvist et al., 2018)

However, a key question remains: How can these gamma-beta interactions, which reflect the combined activity of millions of neurons, be selective enough to control the contents of individual items in WM? WM control, after all, is more than just turning WM “on” and “off”. It is also operations on the individual items held in WM.

We propose that this selective control comes from utilizing network space. In this *spatial computing* paradigm, moving the representation of a WM item from one part of the network to another is considered a computation. Control comes from *where* in network space a specific WM item is held. This allows the item to be accessed and operated on just by knowing its place in network space. Importantly, it enables control without having to know the precise network connectivity forming the ensemble for that item. In this view, spatial computing is mediated by a low-dimensional pattern of gamma-beta power across a network. The control operations on WM contents are reflected in the shared, low-dimensional components of neural activity (Kobak et al., 2019). Spiking carrying the items held in WM is a high-dimensional component that appears where gamma is high and beta is low at each moment in time. Applying a set of WM operations (e.g. executing task rules) corresponds to imposing a low-dimensional spatio-temporal pattern.

To test this idea we compared beta, gamma and spiking activity in several WM tasks where the order sequence of visual objects needed to be retained. Spatial computing predicts that control operations can be read from a low-dimensional spatio-temporal pattern of gamma and beta since this information is spatially organized. The control operations are the task demands. They include the ordering of the objects and their different uses (encoding/maintenance vs determining whether they match a test object). By contrast, gamma and beta are not expected to carry the identity of the items due to their coarser spatial scale than individual neurons. This information should instead be reflected in the high-dimensional spiking because it arises from the connectivity of single neurons, organized on a much finer spatial scale. Further, because the low-dimensional activity reflects WM operations not content, it should have a stable spatial pattern across different sessions where different sets of objects were used. In spiking, we should observe a mix of high-dimensional activity reflecting item identity and the low-dimensional components that account for the current operation being performed on them.

We found support for these predictions and thus offer a novel, spatial perspective on low dimensional activity and the role of oscillations in WM control. We discuss how spatial computing offers a new perspective on mixed selectivity, neural subspaces and redundancy in cortex. Importantly, we discuss how spatial computing provides an account for the generality of WM. In contrast to many computational models of WM, spatial computing allows items novel to a specific task to be operated on without having to re-train networks to the new items.

## Results

We analyzed multiple-electrode neurophysiological recordings from the prefrontal cortex (PFC) of five rhesus monkeys performing four WM tasks. In three of the tasks the monkeys had to remember sequences of objects or colored squares. One task was a single-object delayed match-to-sample task.

We tested the following predictions of spatial computing: 1. There are different neural sources for control-related activity (i.e., top-down task demands) vs the specific items held in WM. 2. The control-related activity is organized as spatial patterns. 3. The patterns are stable across different recording sessions that use different sets of WM items but have the same top-down demands. 4. The control-related activity modulates the spiking of individual neurons such that their activity reflects both the identity of the item held in WM and current task demands.

### Prediction 1: Independent neural sources of control-related and item-specific WM activity

To test this prediction, we used LFP and single neuron activity recorded with acute electrodes in PFC of rhesus monkeys performing a sequence two-item WM task. The monkeys had to remember the identity and order of two visual objects presented in sequence (Task 1, Figure 2A). After a brief memory delay, there was a two-object test sequence that could either be identical to the encoded sample sequence or a mismatch (where either the temporal order or the identity of the objects was changed). After the second test object, monkeys indicated whether the test object sequence matched that of the sample object sequence seen at the start of the trial.

Both item-specific activity (carrying information about each object’s identity) as well as control-related activity (i.e., the requirement to remember the order and to determine if they match a test sequence) are needed to solve the task (Figure 1A, Warden and Miller, 2010). Our previous work demonstrated a ramp-up of spiking (and gamma bursting) at the end of a memory delay that reflected a “read-out” of an item from WM (Lundqvist et al., 2016, 2018). On a population level the ramp up in spiking was selective to order information. Out of the two items held in WM, spiking only increased for the item that was relevant for the upcoming test (information about Sample 1 item before Test 1, information about Sample 2 item before Test 2 in Figure 1. Warden and Miller, 2010; Lundqvist et al., 2018).

**Figure 1.**
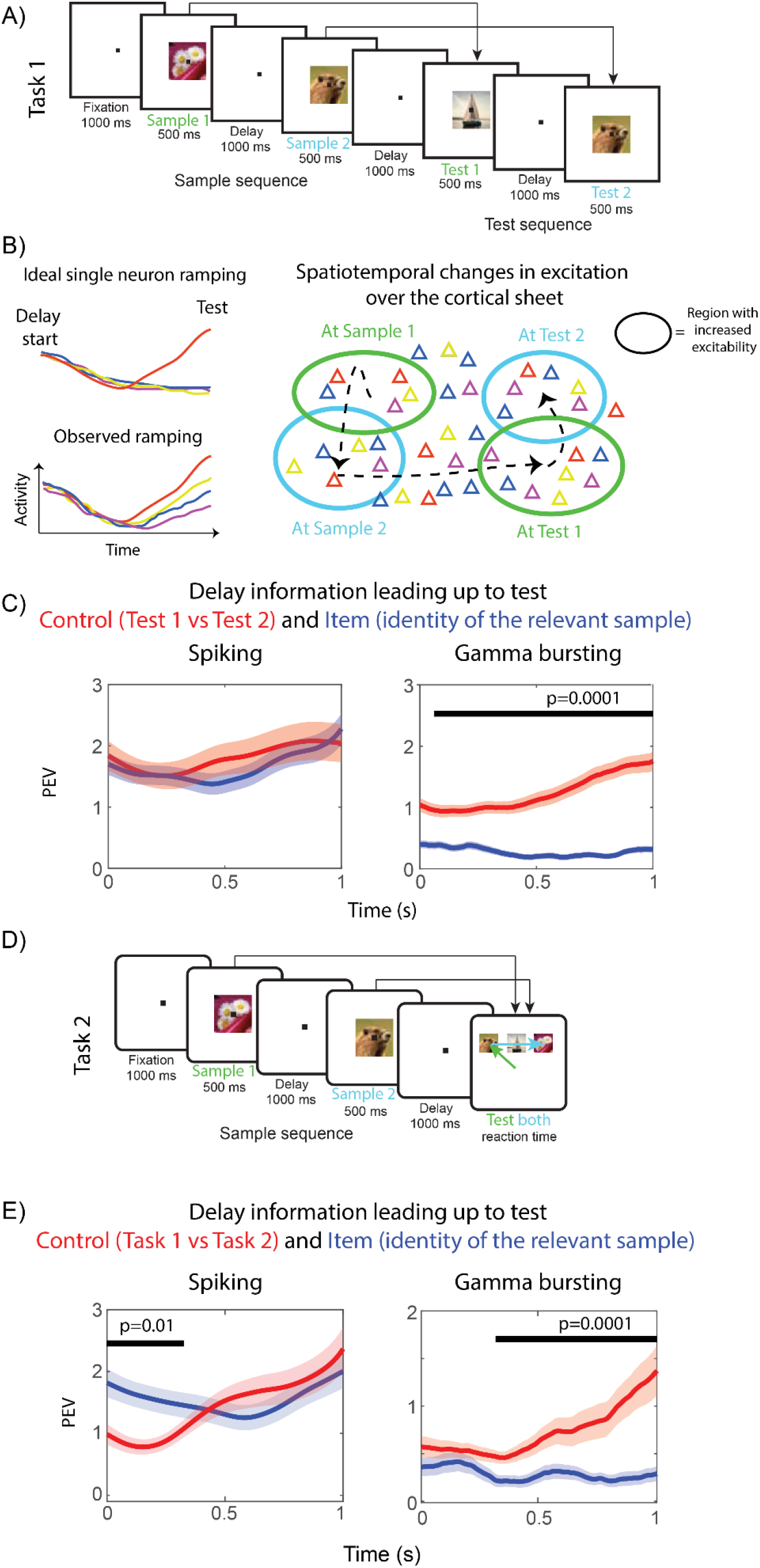
Disassociation between gamma and spiking. A) In Task 1, two object cues are presented and then tested sequentially. In “match trials”, to which monkeys had to respond, the order and identity of the objects have to be the same for sample and test sequences. B) Left: Item-specific activity ramps up before the information about item identity is used in the task. In the idealized case of stimulus selective neurons, their activity only ramps up for one specific cued item. However, recorded neurons with ramping activity respond to all cued items to varying levels (see also Figure S1). Right: the model explains the observed phenomenon of post stimulus ramping after all stimuli to varying degrees by external time-varying excitation that travels over the cortical sheet. Neurons preferring for instance the red or blue cues are scattered throughout the network. Some sites exhibit ramping excitation, reflected in increased activity of all neurons at that site, though to the lesser degree when the preferred object was not provided. C) Red plots show PEV accounting for test order effects estimated over two groups of trial periods, delay periods preceding either Test 1 or Test 2. Blue curves reflect PEV wrt. cued item identity (information about identity of Sample 1 prior to Test 1 and Sample 2 leading up to Test 2 stimuli). Black bars demonstrate when blue and red plots differ, using cluster based statistics. D) Task 2 structure, which is identical to Task 1 until Test 1. Unlike in Task 1 however, at Test 1 both Sample 1 and Sample 2 items are tested in parallel by monkeys making eye-movements to the two targets. E) PEV information pooled over Task 1 and 2 in the delay period leading up to Test 1. Red plots show PEV estimated over periods grouped based on task (Task 1 vs 2), whereas blue curves illustrate PEV information about the cued item identity (average of information about Sample 1 and Sample 2).

This selective ramp-up we observed on the population level was reflected in item-selective single neurons that only ramped up to one of the two tests (demonstrating order specificity). However, when they ramped before a specific test, they did so regardless if their preferred item had been presented during the corresponding sample or not (conceptually illustrated in Figure 1B, single neuron examples shown in Figure S1). An exclusively item-specific ramping neuron would selectively ramp up spiking only for its preferred stimulus, not for others (Figure 1B, left). The observed spiking behaviour could be explained by broad stimulus tuning. Spatial computing suggests that this effect is due to independent origins of control-related and item-specific activity. The ramping comes from top-down control-related excitation that targets cortical locations at much coarser spatial scales than individual neurons. As a result, millions of neurons in these cortical locations receive excitation regardless of their item preference (Figure 1B right). Due to the spatial integration of LFP activity, item-specific activity should largely cancel out in gamma (and beta) bursting. Instead, spatial computing predicts that these LFP oscillations should capture spatially organized information about the order of items but not their identities per se. Importantly, spiking, which probes network activity at much finer spatial scales, should account for both item-specific activity arising from recurrent connectivity and control-related activity inherited from the gamma and beta interactions.

We thus examined spiking and gamma bursting in the delay periods of 1000 ms leading up to the first and second test stimuli (Figure 1A). The data in these periods were labelled by the control-related context (Test 1 or Test 2) and by item identity (i.e. the item that was expected based on the sequence held in WM - the first item from sample sequence for Test 1 and the second item for Test 2). We then calculated the percentage of variance (PEV) in the neural activity explained by these labels. As predicted, the ramp-up of gamma bursting carried information mainly about the control component, order, i.e., whether it was the lead up to Test 1 or Test 2 cue, and not the identity of the retained items (Figure 1C, right panel). Spiking carried a mixture of the two information components (Figure 1C, left), i.e. both the order and the identity of the expected item. We also found a similar difference between spiking and beta activity in line with the prediction about the beta as a correlate of control signals (Figure S2). As expected from the control signal, beta also mainly carried order information.

We further tested this prediction by analyzing data from another WM task (Figure 1A, D). In Task 2, two cues were again sequentially presented, just as in Task 1, but then they differed in how those memories were tested after the memory delay. In Task 2, rather than making a yes/no decision about whether a test sequence matches the remembered one, three objects were presented simultaneously. The monkey had to choose, using eye movements, the correct two objects in the correct order. Monkeys switched back and forth between these two tasks. Note that both tasks are identical up until the testing period after the memory delay. Nonetheless, a previous analysis of this data showed that there were different patterns of spiking ramp-up at the end of the memory delay reflecting the different task demands (Warden and Miller, 2010). Here, we found that control-related information (i.e. whether the monkey was performing Task 1 or Task 2) could be determined from spatial patterns of gamma bursting (Figure 1E, right panel). As predicted by spatial computing, the ramp-up in spiking instead reflected not only the control-related information but also item identity (Figure 1E, left).

We further elaborated on these results by using demixed principal component analysis (dPCA) of the two tasks (Figure 2). dPCA decomposes population activity into principal components dependent on task parameters (Tang et al., 2016). We identified low-dimensional control-related components as well as high-dimensional item-specific components of both gamma bursting and spiking activity. The control-related components captured two types of activity: differences between the two tasks (“Task individual”) and shared patterns of activity over time in the two tasks (“Task general”). Item components reflected the item-specific spiking. As predicted, the control-related components were more dominant in gamma bursting (Figure 2, left) whereas spiking additionally captured strong item dependent components (Figure 2, right panel).

**Figure 2.**
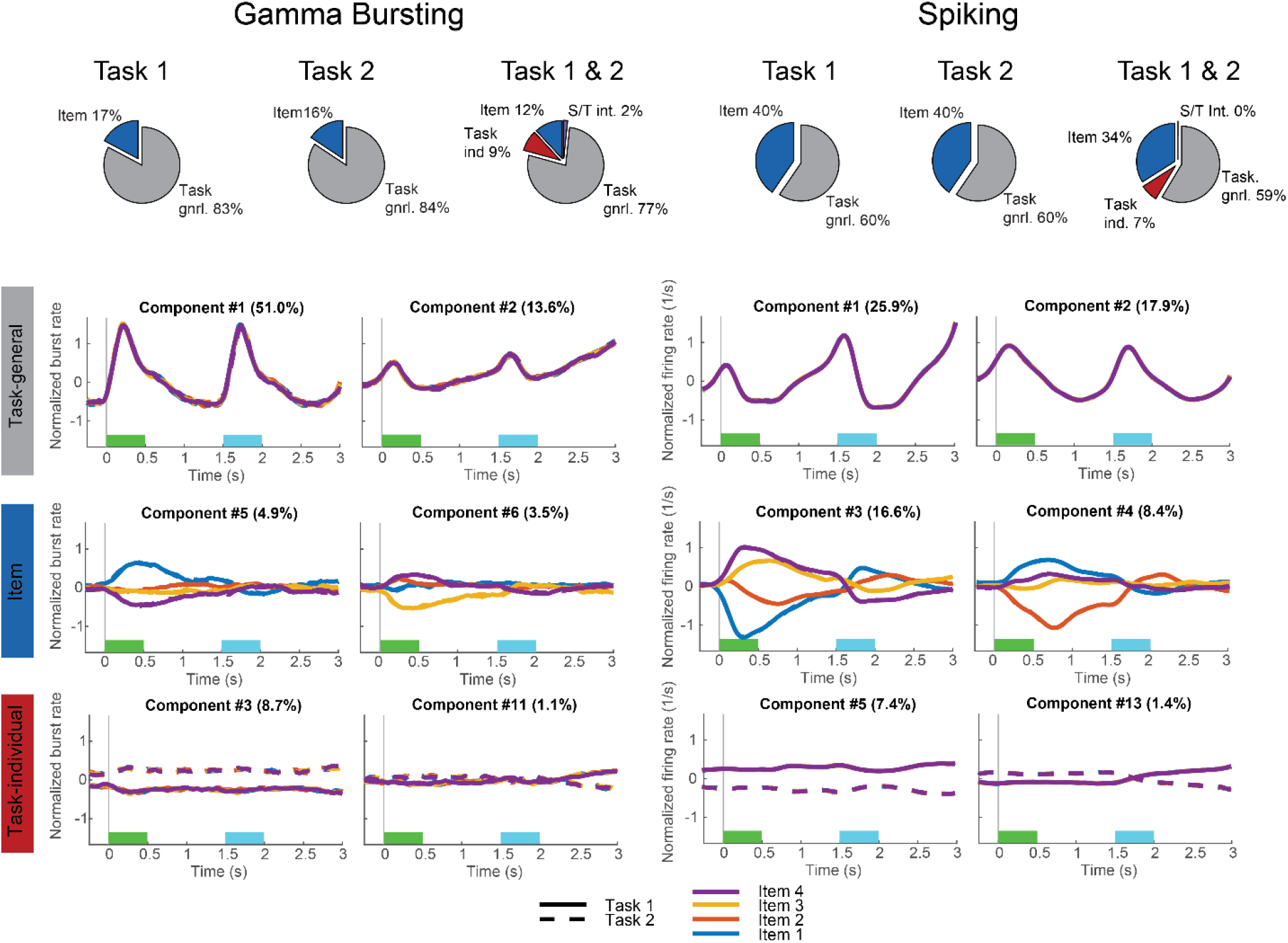
Demixed principal component analysis (dPCA) of gamma and spiking. *Spatial computing implies that task dependent components (red and grey) should be more prevalent in gamma bursting as compared to spiking (c.f. Figure 1). Here we used dPCA (Kobak et al., 2016) to extract the principal components and attribute them to task control-related and item-specific activity. “Task general” (grey) and “Task individual” (red) components correspond to low-dimensional task control-related activity. “Task general” components reflect shared patterns of activity over time in the two tasks whereas “Task individual” components explain the variance that originates from the differences between the two tasks. Item dependent components (blue) account for the variance between four different cued items (item-specific activity)*. The bottom half of the figure shows example components for Task 1 and Task 2 combined.

One possibility is that the observed difference between gamma and spiking simply reflected differences in the quality in these measures of cortical activity (spikes vs LFPs). In other words item-specific information might simply be more difficult to read out from gamma. To test for this, we performed the same analysis on another WM dataset referred to as Task 3 (Figure 3A, Lundqvist et al., 2016). As in the two previous tasks the items were presented sequentially. Though this time each sample was in a different extrafoveal location instead of all foveally as in Task 1 and Task 2. As a result, we expected the item-specific spiking to be spatially distributed due to retinotopic organization. Therefore spatial computing predicts that we should not observe the same disassociation between control-related and item-specific activity in spiking and gamma, as in the previous tasks. Both types of information should be spatially organized, meaning that gamma and spiking should now carry similar contents. The results shown in Figure S3 are consistent with the prediction. There was similar amount of control-related and item-specific activity in both neural measures. This suggests that differences between the gamma and spiking reported in Tasks 1 and Task 2 reflected distinct spatial organization of control-related and item-specific information. It is not just that gamma is merely a low-pass filtered spiking activity. Further, in line with our earlier discussion of gamma-beta interactions as spatial computing correlates, we found that beta bursting was dominated by the low-dimensional, task dependent components as well (Figure S3).

**Figure 3.**
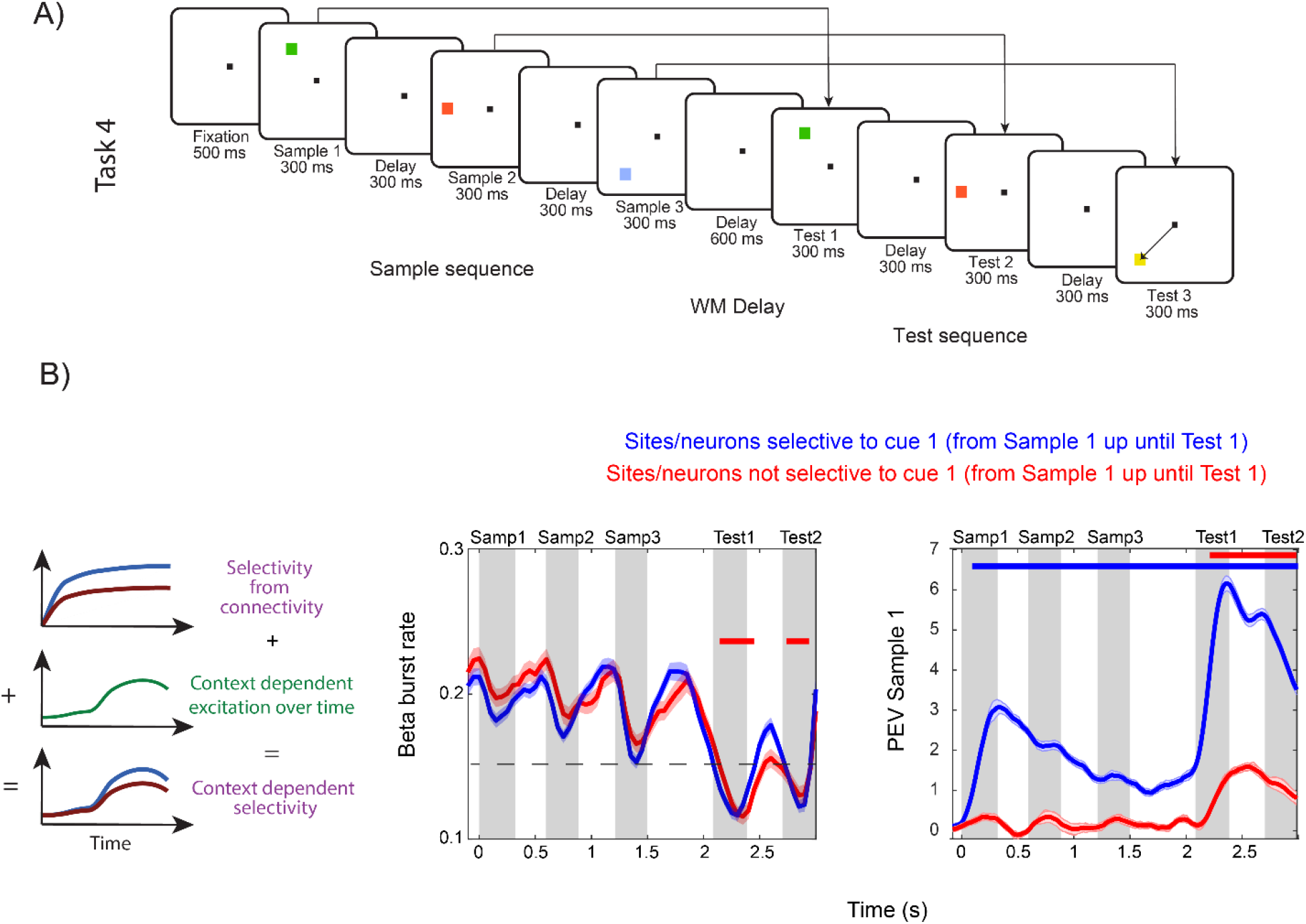
Control-related (task demand dependent) selectivity arising from time-varying excitation. A) In Task 3, two or three colored squares are presented in a sequence. Their location and color are to be remembered. Following the 0.6 s delay, the sequence of squares is repeated. Monkeys should saccade to the square in the test sequence that changed color relative the sample sequence. B) Left: the conceptual illustration how in the proposed spatial computing paradigm mixed selectivity arises from a combination of item selectivity (given by network connectivity) and the unique pattern of local excitation (local footprint of a spatially distributed pattern of excitation hypothesized to control WM). Middle: beta bursting reflects (is anti-correlated with) the excitation in the network. Blue curves correspond to sites in which neurons are selective to items cued at Samples 1-3, red curves describe sites with no such selective neurons. Trials with three sequential sample cues and two test cues (monkeys correctly respond at Test 2) are shown. Dotted line corresponds to the lowest beta burst rate for selective sites when not including the Test periods. Red bar denotes times where beta burst rate in non-selective sites drops below that rate (cluster based statistic, p<0.05). Right: PEV in spiking accounting for the information about an item presented at Sample 1 measured in neurons on sites that are item selective (blue) and non-selective (red) during sample and delay periods.

### Predictions 2 and 3: Control-related activity form spatial patterns that are stable across sessions

Spatial computing predicts that control-related spiking and LFP activity is spatially distributed and can be decomposed into spatial components reflecting patterns of excitation. To test this hypothesis, we used data from dense chronically implanted electrode arrays, which enabled us to 1) map the spatial distribution of various components, and 2) assess their stability across different recording sessions. We recorded simultaneously from 4 Utah arrays, implanted in left and right PFC. The monkeys performed an object delayed match-to-sample task over multiple sessions (Task 4, Figure 4). Different sets of 8 objects (WM items) were used for each session.

**Figure 4.**
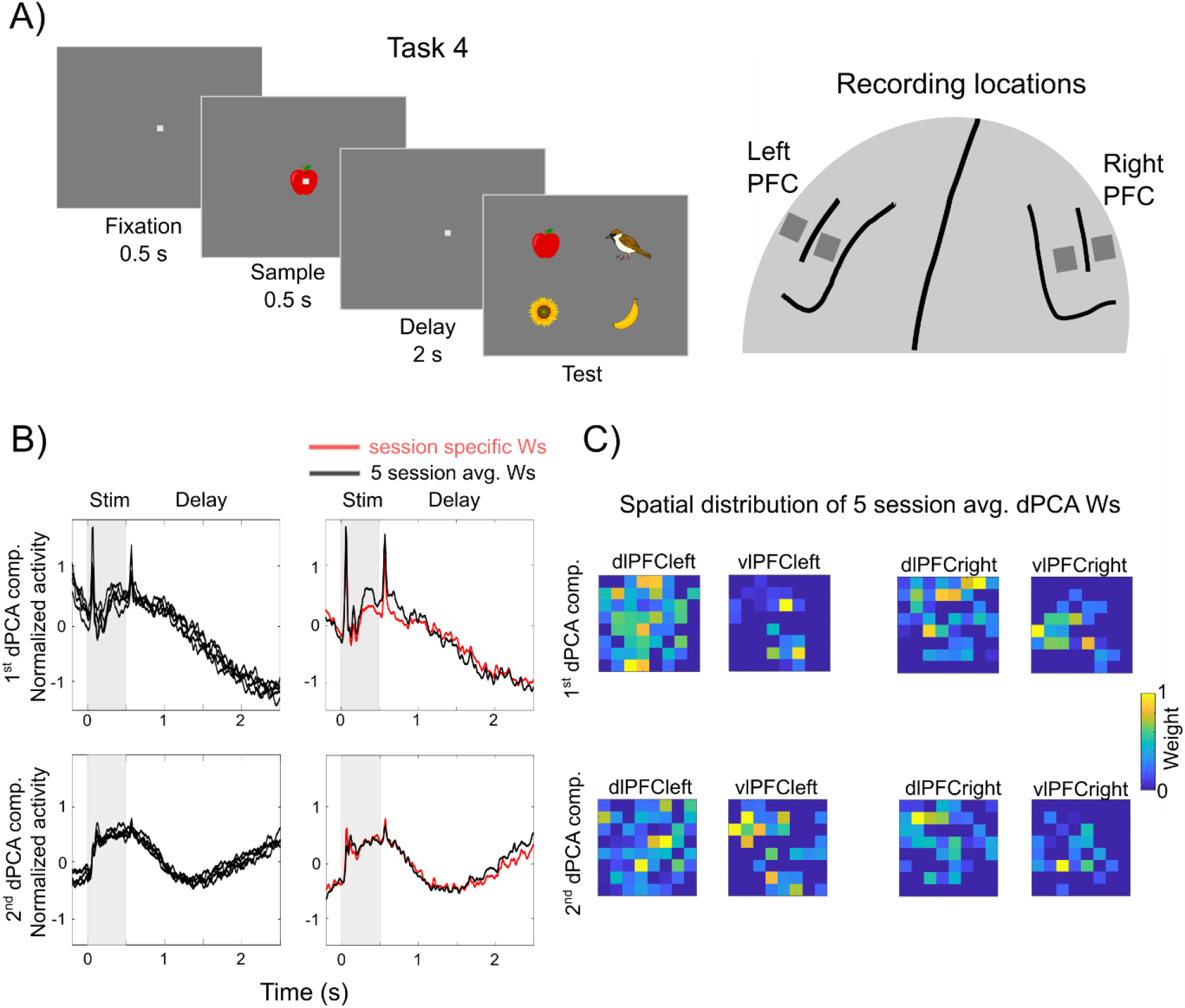
Spatial patterns of control-related activity (reflecting task demands). A) Left: Schematic depiction of the delayed match-to-sample task. Right: schematic depiction of the recording locations of the four chronically implanted Utah arrays (left and right dlPFC, left and right vlPFC). B) Left: the 1st (top) and 2nd (bottom) task-general dPCA components of gamma activity projected using the five-session average dPCA weight vectors for each one of the five sessions. Right: plotted are the five-session averages of the 1st and 2nd task-general dPCA gamma components using i) the five-session average dPCA weight vectors (black) and ii) session-specific weight vectors (red). C) The 4-array spatial distribution of the session-average normalized dPCA weights used in B) for the first (top) and second (bottom) task dependent dPCA components.

We focused on gamma activity since for each electrode it reflects the activity from a stable, spatially located group of neurons across sessions (whereas the same individual single neurons may not remain isolated across sessions). We performed dPCA analysis of the gamma activity (45-100 Hz) within the delay period in each session. We focused on the first two control-related components that reflected shared activity over time regardless of item identity. They explained on average 75% and 8% of the data variance, respectively. To spatially localize the two components in each recording session, we extracted their corresponding dPCA loadings (weights used for projecting the data). Further, to minimize the risk of overfitting the dPCA model to session specific noise we sparsified the weights. We only kept those weights that significantly contributed to the component (see Materials and Methods). We then averaged the dPCA weights across five sessions per component and used the two resulting weight vectors to derive the corresponding projections for each session. This yielded virtually identical components in each session (Figure 4B, left). Further, session-specific dPCA components, i.e. based on loadings extracted from individual sessions, showed striking resemblance to those obtained with the five-session averaged weight vector (Figure 4B, right). Thus the spatial distribution of these components was stable across sessions (and across sets of WM items). We plot their spatial organisation in Figure 4C to illustrate that they were distributed in all four recorded areas and distinct for the two components, as predicted by spatial computing.

The difference between the two control-related components is also reflected over time. The first component corresponded to activity that slowly decayed over the delay period. The second one corresponded to activity that initially decayed but then ramped up before the end of the delay. The components also exhibited differences during the sample period. The first component was elevated throughout the sample. The second component had activity primarily at the onset and offset of sample. Since the components were selected solely based on the delay activity, these differences in other task periods implies that they detected consistent differences between recording locations (and not over-fitting to data).

### Prediction 4: Low-dimensional spatial excitation gives rise to context dependent selectivity

One of the implications of spatial computing is that at the single-neuron level, there will be a mix of control and item-specific activity. This arises out of control-related, low-dimensional spatial patterns of excitation combining with the item-selectivity of individual neurons. To test this hypothesis, we analysed spiking and LFPs from the third WM task (Task 3, Figure 3A). According to the spatial computing paradigm, changes in excitation levels should increase or decrease item-selectivity of single neurons depending on current task demands (e.g. different task contexts like encoding of items into WM vs testing those memories at the end of the trial). We considered here beta and gamma bursting as proxies for local excitation. In particular, beta bursts should reflect excitation levels distinguishable from spiking. This is because they express less cross-talk between spiking and field activity than gamma (Ray and Maunsell, 2011). We have previously shown that beta bursts are anti-correlated with bursts of gamma and spiking (Lundqvist et al., 2016; 2018). For example, beta bursting is lower when the sample objects are presented and gamma bursting is at the same time elevated (Figure 3B; Figure S5).

Here, we found that the levels of beta activity had a distinct shift downwards before and during test periods compared to sample encoding periods (Figure 3B) suggesting a general shift in excitability. It was not due to saccades as it was seen equally during test periods in which animals responded or withheld the response (compare Test 1 (no response) and Test 2 (response) in Figure 3B). This implies that excitation levels indexed by LFPs (and approximated here by beta) were generally elevated during test periods compared to encoding. In line with earlier analysis of the same data (Lundqvist et al., 2016), sites with neurons item-selective activity had lower beta during encoding (sample cues) than sites with non-selective neurons (Figure 3B; using units selective to Sample 1 in the first 600 ms after the sample onset, p<0.05, cluster based statistic). However, during the test periods beta for non-selective sites was consistently lower compared to the lowest beta observed at selective sites during encoding. This more widespread excitation (reflected in LFPs) should according to the hypothesis be manifested in a more widespread item-selectivity (of single neurons) in the network. In trials where the same items were presented during Sample 1 and Test 1, beta was suppressed (Figure 3B), and gamma power (Figure S5) and firing rates were increased during the test relative the sample period. Importantly, item-selective activity was confined to a smaller part of the network during the encoding compared to the test periods (Figure 3B, right). A significantly larger portion of the neurons responded to Test 1 than Sample 1 stimuli, even in trials in which they were identical (62/496 had significant variance explained by Sample 1 identity compared to 152/496 by Test 1 identity using the first 600 ms from the onset of sample cue and test cue, respectively; p<0.05, cluster based statistic). This suggests that controlling the spatio-temporal patterns of excitation levels could be an effective way to distinguish information in the selective neural population between the sample and test periods, as a natural implication of the spatial computing paradigm.

## Discussion

We have tested the hypothesis that the PFC implements *spatial computing* for the control of WM. The central idea is that top-down control-related information (e.g., the ordering of multiple items held in WM, how they are used etc) operates in network space. Different WM items are distributed across different locations in a network so they can be manipulated independently. The manipulations are reflected in neural oscillations that act on hundreds of thousands of neurons (rather than target single neurons) at different network locations. The manifestation is a dynamic patchwork of beta and gamma bursts. As in our earlier framework, the gamma bursts co-register with the spiking carrying the individual items held in WM. Beta bursts act as an inhibitory control signal that inhibits gamma/spiking when and where beta power is high (Lundqvist et al., 2016; 2018; Miller et al., 2018).

To understand how this works, consider our task requirement to retain two objects (A and B) in the order in which they appeared (1^st^ or 2^nd^). The idea is that during the initial encoding of the sequence, different patterns of gamma are activated for the first vs second item. The gamma patterns are determined by patterns in beta (the top-down control signal): Gamma is where beta is not. When the first object is shown (say, object A), the underlying network has a unique gamma pattern corresponding to “1^st^ item”. This injects excitation and primes neurons that are selective to A in those gamma patches. When the second object (say, B) is shown, top-down control will form a different gamma patch pattern corresponding to “2^nd^ item”. Neurons selective for B are then excited and primed in those patches. To read out which object was first (or second), the corresponding “read-out” gamma pattern is activated. The primed neurons from that patch (network location) will spike more strongly. The concept rests on the assumption that there is significant redundancy in cortex such that there are neurons selective to both A and B in both patterns of patches (corresponding to 1^st^ and 2^nd^ object; Goldman-Rakic, 1996). In short, spatial computing posits that WM control stems from spatio-temporal activity patterns across network space that reflect and change with top-down task demands.

This separation of control (in this case, order) from content (A vs B) endows WM the ability to generalize and the transfer learning to new items not used when a given task was initially learned, i.e., zero-shot learning (O’Reilly and Frank, 2006). This can be explained by different sources for these signals. Cortical beta oscillations are thought to emerge from loops between thalamus, cortex and subcortical structures (Ketz et al., 2015). This suggests that top-down information is imposed, at least in part, from outside the local PFC cortical network itself (O’Reilly and Frank, 2006; Schmitt et al, 2017; Lusk et al., 2020). The top-down excitation would selectively support stable retention of information in those parts of the network it targets. In support of this view, there is growing evidence suggesting that excitation from mediodorsal thalamus is needed to sustain working memory and attention activity in PFC (Bolkan et al., 2017; Schmitt et al., 2017; Lusk et al., 2020). The selectivity of individual neurons for specific items, by contrast, may be more dependent on patterns of recurrent connectivity and inputs local to cortex (Goldman-Rakic, 1996; Ko et al., 2013; Kritzer et al., 1996; Rao et al., 1999).

We found evidence supporting the spatial computing hypothesis. The spatial pattern of beta and gamma reflected control operations of the task at hand. This included item order, how each item was currently being used (encoding/maintenance vs ‘reading out” the items to make a match judgement), and which of two WM tasks the monkey was performing. Notably, the two tasks were identical except for how WM items would be read out and used at the end of the trial (recognizing a match vs choosing matches from a number of alternatives). Consistent with the spatial encoding of control signals, control-related information was found in the pattern of gamma/beta activity, shared among neurons within a few hundred micrometers. Consequently, it arose on a larger spatial scale than the item-specific activity that had more of a salt-and-pepper, finer grained, distribution. This directly implies that the spatial dimensions are used and that oscillations can be used to selectively control information despite acting on millions of neurons simultaneously. Importantly, we observed that gamma and spiking, though highly correlated in time and space, carried different information. The gamma pattern reflected top-down control information per se. By contrast, spiking carried information about specific WM items as well as top-down information. The top-down information was “inherited” by virtue of which gamma patch each neuron belonged to.

By examining recordings from chronically implanted electrode arrays, we also found that the low-dimensional gamma and beta patterns were consistent across recording sessions. Within a session, they were dynamic, changing with current task demands. Nonetheless, these spatio-temporal dynamics were similar across multiple recording sessions that used different WM items. We previously reported that a ramp-up of gamma bursting was related to read-out from WM (Lundqvist et al., 2016; 2018). Here, we showed that the same parts of the network (recording array) showed gamma bursting ramp-up even though those sessions used different WM items. Spiking during these ramp-ups carry information about the specific item being read-out (Lundqvist et al., 2016; 2018). Thus, our current results suggest that the same network locations in the PFC are used for the same operation (read-out) regardless of item identity.

Spatial computing is consistent with, and sheds new light on, a variety of observations in the extant literature. Spatial computing requires representational redundancy. Information about a given WM item is represented in multiple parts of a network (for different operations). Such redundancy seems to be a hallmark of cortical activity (Pasternak and Greenlee, 2005; Siegel et al., 2015; Dotson et al., 2018; Stringer et al., 2019). It could be supported by horizontal connections between neurons with shared stimulus preference (Goldman-Rakic, 1996; Kisvarday et al., 1997; Ko et al., 2013). Spatial computing is also in line with growing interest in the dimensionality of cortical representations (Rigotti et al., 2013; Stringer et al., 2019; Badre et al., 2020; Cueva et al., 2019; Wolff et al., 2020; Kobak et al., 2016; Tang et al., 2019; MacDowell et al., 2020). Low-dimensional activity, i.e. shared across many neurons and across experimental conditions (such as our patterns of gamma and beta patches) has been implicated in the ability to generalize across tasks (Badre et al., 2020). Indeed, low-dimensional activity often reflects the structure of tasks, e.g., modulation of activity across different task periods (Tang et al., 2019). Changes in low-dimensional activity have been shown to correlate with task learning and reflect time within a trial (Tang et al., 2019; Cueva et al., 2019; Wolff et al., 2020). Our findings add that these arise from spatial patterns of excitation. Spatial computing is also consistent with observations that population spiking accounting for WM items rotate into distinct subspaces depending on whether the item is currently behaviorally relevant (Panichello and Buschman, 2021). Spatial computing suggests that these rotations are driven by the spatial flow of information in networks. Moving the information would cause a rotation into a new subspace in this view.

Spatial computing can also explain classic observations that the spiking of individual neurons is highly task-dependent (Goldman-Rakic et al 1996). Some neurons, for example, only respond to an item when it is a to-be-remembered sample or only when it is a test item used to compare against the memorised content. Spatial computing explains this by having different sets of gamma patterns being activated for these different task contexts (see Figure 3, Figure 4). The neurons only spike to a preferred item when its gamma patch is active. Similar results have been found in models of artificial recurrent neural networks trained on WM tasks. The artificial network units form multiple functional neuronal sub-groups with some units activating to an item when it was a sample cue, others – when it was a test item (Dubreuil et al., 2020). Here we found similar sub-groups experimentally with the addition that they were spatially organized. This, in turn, may also provide insights into mixed selectivity (Rigotti et al., 2013; Fusi et al., 2016). Neurons with mixed selectivity show context-dependent spiking that is non-linear. It cannot be predicted from their responses to the individual elements that combine to make that context. Mixed selectivity is thought to add computational horsepower, increase network storage capacity among other functional benefits (Rigotti et 2013; Fusi et al, 2016). Here, we add that the context-dependent activity of mixed selectivity neurons arises from the top-down control low-dimensional patterns applied to networks, not just from the detailed connectivity within the network. Thus the computational benefits of mixed selectivity may be flexibly adapted from task to task.

In sum, spatial computing postulates a novel mechanism for how neural oscillations may implement selective WM control. It can also explain how PFC networks may consequently achieve flexible WM with powerful generalization capabilities. In doing so, it offers a new perspective on the functional role of low-dimensional activity that often seems to dominate cortical activity.

## Materials and Methods

### Data from previous studies

We analyzed data from two previous studies (Warden and Miller, 2010; Lundqvist et al., 2016). In total the two studies included three experimental tasks (Task 1 & 2 from Warden and Miller, 2010; Task 3 from Lundqvist et al., 2016). For details of training and data collection, please see those studies. Briefly, each task involved two Rhesus macaques that were trained until they performed well above chance. They were trained with positive reward (juice) only and maintained in accordance with the National Institutes of Health guidelines and the policies of the Massachusetts Institute of Technology Committee for Animal Care).

For each recording, a new set of acute electrode pairs (tungsten, epoxy-coated, FHC) was lowered through a grid. Between 8 and 20 prefrontal electrodes were recorded from simultaneously on each session (34 sessions for Task 1 and 2, 30 sessions for Task 3). Task 1 and Task 2 were recorded during the same sessions in a blocked design. Only electrodes containing isolatable units were kept for further analysis.

### Data recordings

We recorded data from one Rhesus macaque monkey performing a delayed-match-to sample task (Task 4). It was trained with positive reward (juice) and maintained in accordance with the National Institutes of Health guidelines and the policies of the Massachusetts Institute of Technology Committee for Animal Care). It had 4 Utah arrays (64 channels each) chronically implanted in left vlPFC, left dlPFC, right vlPFC and right dlPFC. We recorded at a sampling rate of 30 kHz. We recorded from 5 sessions. There were 8 possible objects to be held in working memory each sessions, and 3/8 of the remaining objects acted as distractor at test. Sessions 2 and 3 had the same set of possible objects, and sessions 4 and 5 also shared the same set, otherwise there was no overlap across sessions (in other words, there were 3 unique sets). Over the 5 sessions the monkey performed at a high accuracy (94%), not including fixation breaks. We only analyzed the correct trials (2758, 2526, 2413, 2110 and 2306 in the 5 sessions).

### Signal Processing

#### Preprocessing Task 1 and 2

At first, all electrodes without any isolatable neurons were removed. Then, a notch filter with constant phase across a session was applied to remove 60-Hz line noise and its second harmonic. On some sessions there were high-power, broadband frequency artifacts; these sessions were discarded from further analysis.

#### Preprocessing Task 3

We first removed apparent noise sources from the signal. In particular, a notch filter was applied to remove 60-Hz line noise with constant phase across a session. In addition, we removed periodic deflections seen in the evoked potentials (every 47 ms, lasting 1 ms, on a subset of electrodes, phase locked to stimulus onset). The signal was filtered and down sampled to 1 kHz (from 30 kHz).

#### Preprocessing Task 4

We removed channels that had bad contacts and much higher (noise) amplitudes than the rest using visual inspection. Data was down sampled to 1 kHz and 60 Hz noise removed.

For spectral analysis we applied multi-taper analysis (with a family of orthogonal tapers produced by Slepian functions; Slepian, 1978; Thomson, 1982; Jarvis and Mitra, 2001). The multi-taper approach was adopted with frequency-dependent window lengths corresponding to six to eight oscillatory cycles and frequency smoothing corresponding to 0.2–0.3 of the central frequency, f0, i.e., f0 ± 0.2f0, where f0 were sampled with the resolution of 1 Hz (this configuration implies that two to three tapers were used). The spectrograms were estimated with the temporal resolution of 1 ms. Typically we present total power of raw LFPs (after removal of noise) without subtracting any baseline or estimated evoked content.

### Burst Extraction

To extract bursts of high-power events on a single trial level we utilized a previously developed method (Lundqvist et al., 2016; 2018a). In the first step of the oscillatory burst identification, a temporal profile of the LFP spectral content within a frequency band of interest was estimated. We used two alternative methods of spectral quantification (see above). We either narrow-band-filtered LFP trials and extracted the analytic amplitudes (envelope) or we used single-trial spectrograms, obtained with the multi-taper approach, to calculate smooth estimates of time-varying band power (all presented results were obtained with the multi-taper approach; the results for the two methods were very similar). Next we defined oscillatory bursts as intervals during individual trials when the respective measure of instantaneous spectral power exceeded the threshold set as two SDs above the trial mean value for that particular frequency, and with the duration of at least three cycles. Having the burst intervals extracted for the beta band (20–35 Hz) and three gamma sub-band oscillations (40–65, 55–90, and 70–100 Hz) from each trial, we defined a single-trial point process (binary state: no burst vs burst within a 10-ms window) with the resolution of 10 ms and trial-average measure, a so-called burst rate for each spectral band. This quantity corresponds to the chance of a burst occurrence on an individual electrode at a particular time in the trial (a proportion of trials where a given electrode displays burst-like oscillatory dynamics around the time point of interest sliding over the trial length).

### Statistical Methods

The majority of tests performed in this study were nonparametric due to insufficient evidence for model data distributions. To address the multi-comparisons problem, we employed Kruskal-Wallis, Friedman’s, and Wilcoxon’s signed-rank tests where appropriate. In addition, for the comparison between temporal profiles of the normalized firing rates within versus outside oscillatory bursts, we resorted to a permutation test on the largest cluster based statistics (Maris and Oostenveld, 2007), originally proposed to increase the test sensitivity based on the known properties of the data (here being temporal dependency). Finally, some attention should be given to the way we report correlations between the measures of time-varying spectral band content and burst rate statistics. The correlation analyses were performed on individual electrodes and only the summary statistics (mean and SE) for the electrode-wise significant effects (p < 0.01) are presented.

### Estimation of information

The bias-corrected PEV (Olejnik and Algina, 2003) was estimated across trials with different conditions from firing rates averaged in 50-ms bins across trials within each trial. We performed two-way ANOVA where trials had multiple groupings (i.e. stimulus or delay/task). All correct trials were used, as the groups were well balanced each session. The bias correction was used as it avoids the problem of non-zero mean PEV for small sample sizes.

As a result, (bias-corrected) PEV allowed for the quantification of information carried by the modulation of firing rates or burst rates of individual units accounting for the stimulus, task or task period (delay 1 vs delay 2 in Figure 2).

For Figure 4 we used PEV to estimate selective and non-selective units. Here, for a unit to be classified as selective it needed to have a p-value in the ANOVA test for Stimulus 1 < 0.05. We estimated selective units based the on the 300 ms of Sample 1 presentation and the following 300 ms of delay. We did not use the full delay up until test since we wanted to compare the selectivity to that during and following the Test 1 on equal footing (which had 300 ms of test period and a 300 ms delay before Test 2). Another reason to avoid using the full delay was to demonstrate that the addition of selective units in the previously unselective population was not simply due to passage of time but timed to test onset. We used all correct trials in which Sample 1 was equal to Test 1 (to increase the statistical power we used both load 2 and load 3 trials combined since they had the same Sample 1 and Test 1 timings). In these trials the same items were presented in Test 1 and Sample 1, and the monkeys responded neither during Test 1 nor the following delay but instead awaited Test 2.

### Demixed principal component analysis

To identify low-dimensional manifold for neural activity, we performed a demixed principal components analysis (dPCA) (Kobak et al., 2016). This approach allows not only for compressing the data, similarly to PCA, but also separates the underlying components with respect to the requested task parameters by demixing the dependencies of the population activity on the task parameters. In a nutshell, demixing is achieved by minimising the reconstruction error between the projections and the neural activity averaged over trials (unlike in PCA where the reconstruction error on single trials is minimised) and over the requested task parameters. In addition, when compared to PCA the method used here benefits from greater flexibility offered by using two different linear mappings for encoding vs decoding. More technical as well as theoretical details of dPCA can be found in (Kobak et al., 2016).

In our analyses dPCA was applied to both spiking data (firing rates obtained by convolving the spike point process with 50-ms wide Gaussian kernel) and oscillatory bursts in beta and gamma bands (burst point process convolved with 50-ms wide Gaussian kernel). To achieve demixing effect we grouped trials into task- (Task 1 vs Task 2) and stimulus- (Sample 1) dependent sets, and analyzed trials in the interval from 100 ms prior to the first sample cue (Sample 1) until the first test cue (Test 1). Apart from task and stimulus-dependent components, dPCA also produced a condition-independent component corresponding to low-dimensional time-dependent task activity.

We also used dPCA to study spatial distribution of LFP activity patterns. To this end, dPCA loadings (weights) were extracted for the condition-independent components in each session and, to avoid overfitting to noise, a greedy search for the minimal set of weights that preserve the original component, i.e. with the mean square error below the fixed threshold, was employed. As a result, the weight vectors were made sparse by effectively removing the contribution from 42% of electrodes on average (ranging from 34% to 55%). The saliency of the contribution of particular electrodes to each component was attributed to the absolute value of the corresponding loading coefficient in the reduced weight vector.

## Data availability

All preprocessed data and code are available online, relevant raw data will be available from the corresponding author on reasonable request.

## Supplemental Figures

**Figure S1.**
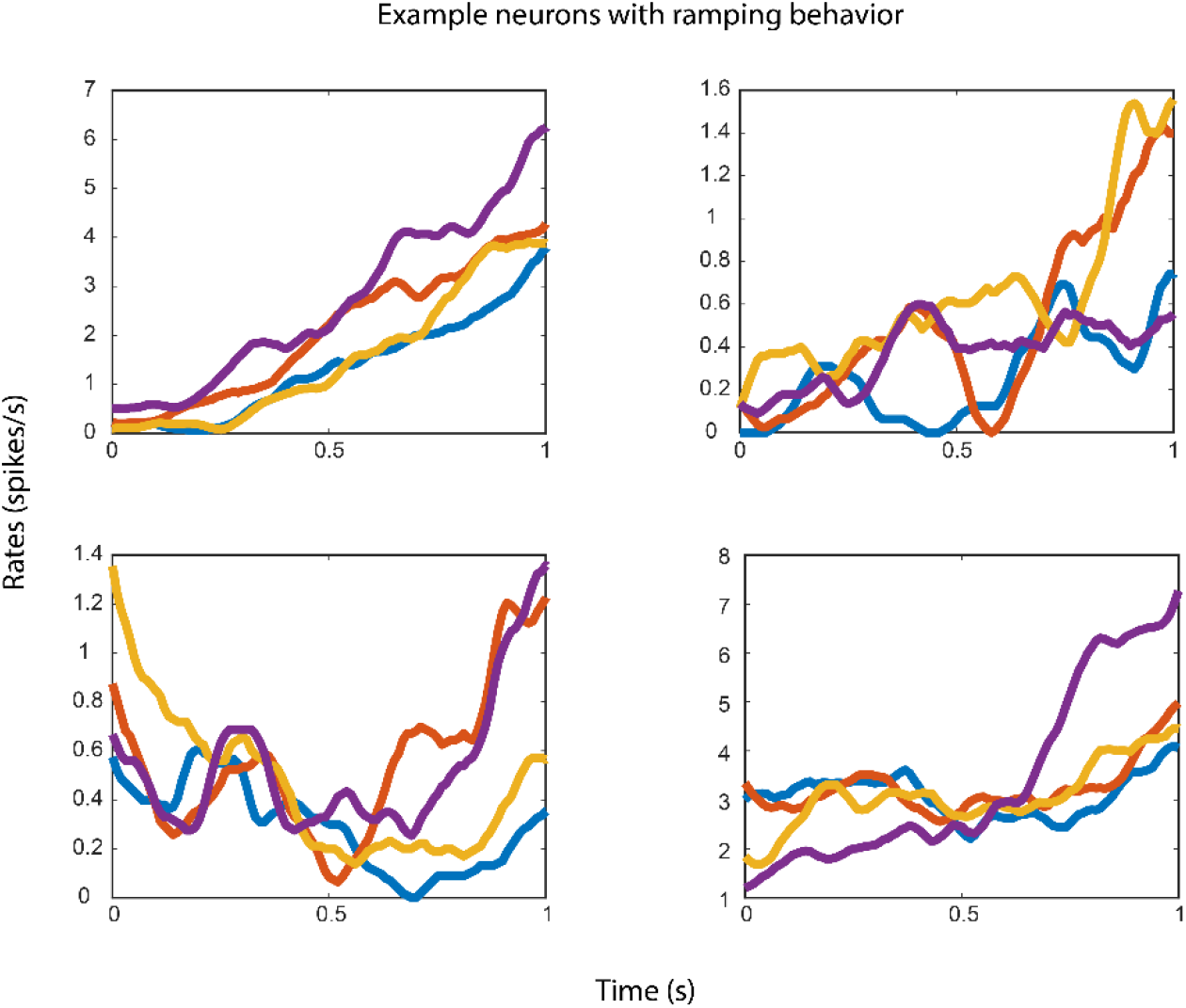
Single neuron examples of ramping activity. Shown are the activity grouped by sample identity (blue/red/yellow/purple) for all four neurons with ramping activity from two random sessions.

**Figure S2.**
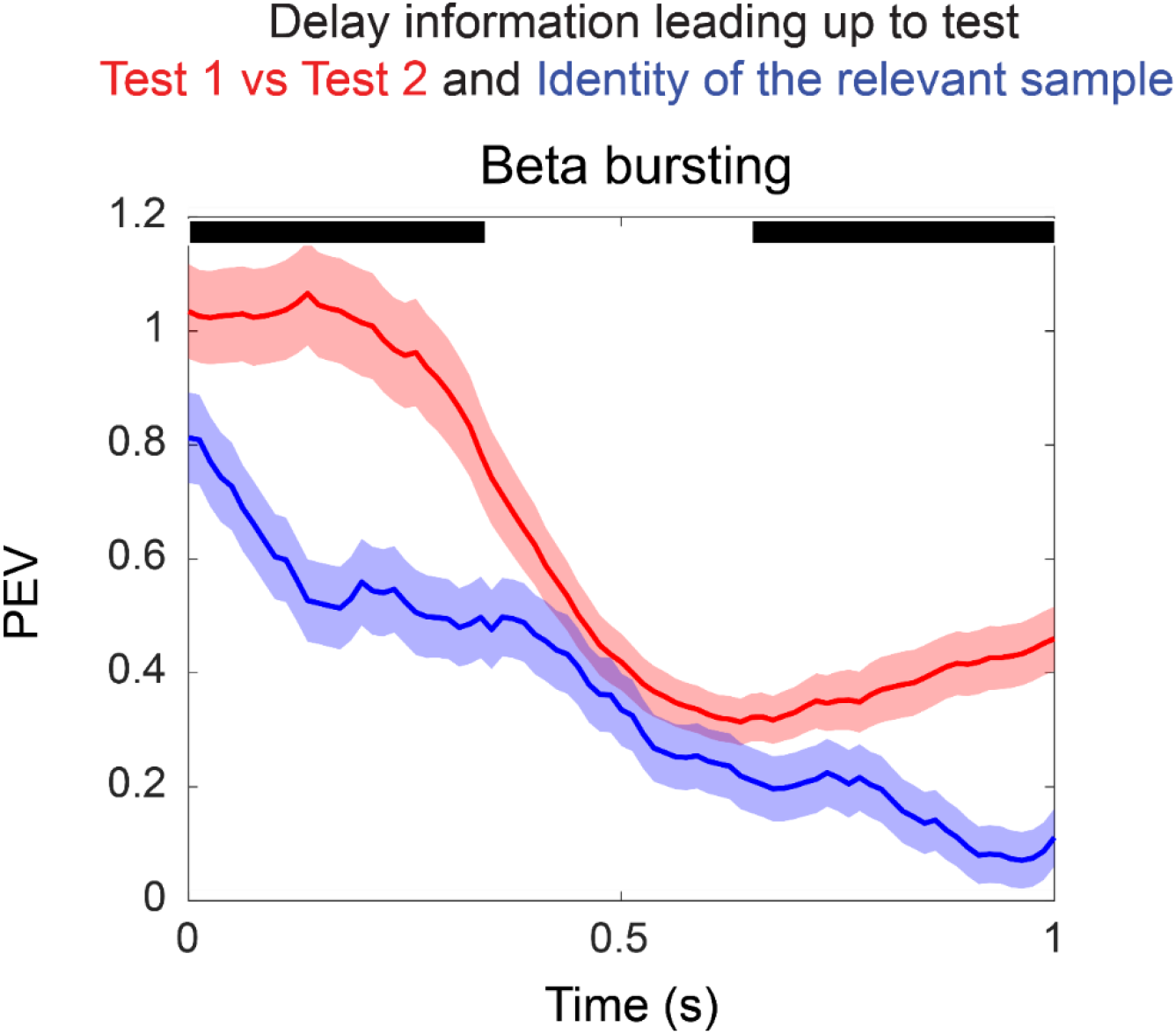
Beta information about sample and upcoming test. Red plots show PEV accounting for test order effects estimated over two groups of periods, preceding either Test 1 or Test 2. Blue curves reflect PEV wrt. sample identity (information about identity about Sample 1 prior to Test 1 and Sample 2 leading up to Test 2). Black bars demonstrate when blue and red plots differ, using cluster based statistics (p<0.05).

**Figure S3.**
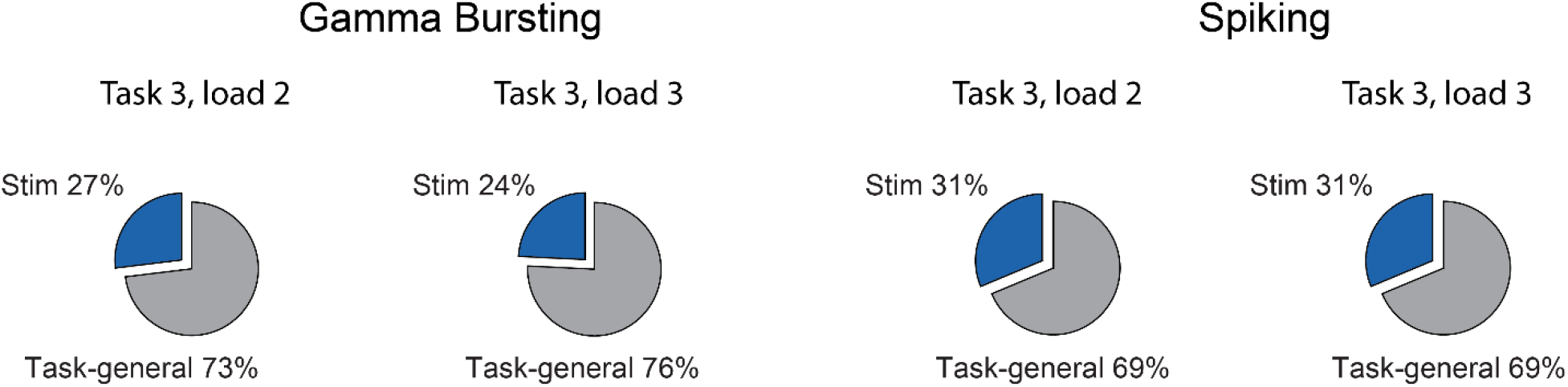
dPCA analysis of Task 3. Shown are the proportion of variance that can be attributed to stimulus (blue) and condition-independent (grey) components for gamma (left) and spiking (right). Due to their distinct task structures, load 2 and load 3 were analyzed separately.

**Figure S4.**
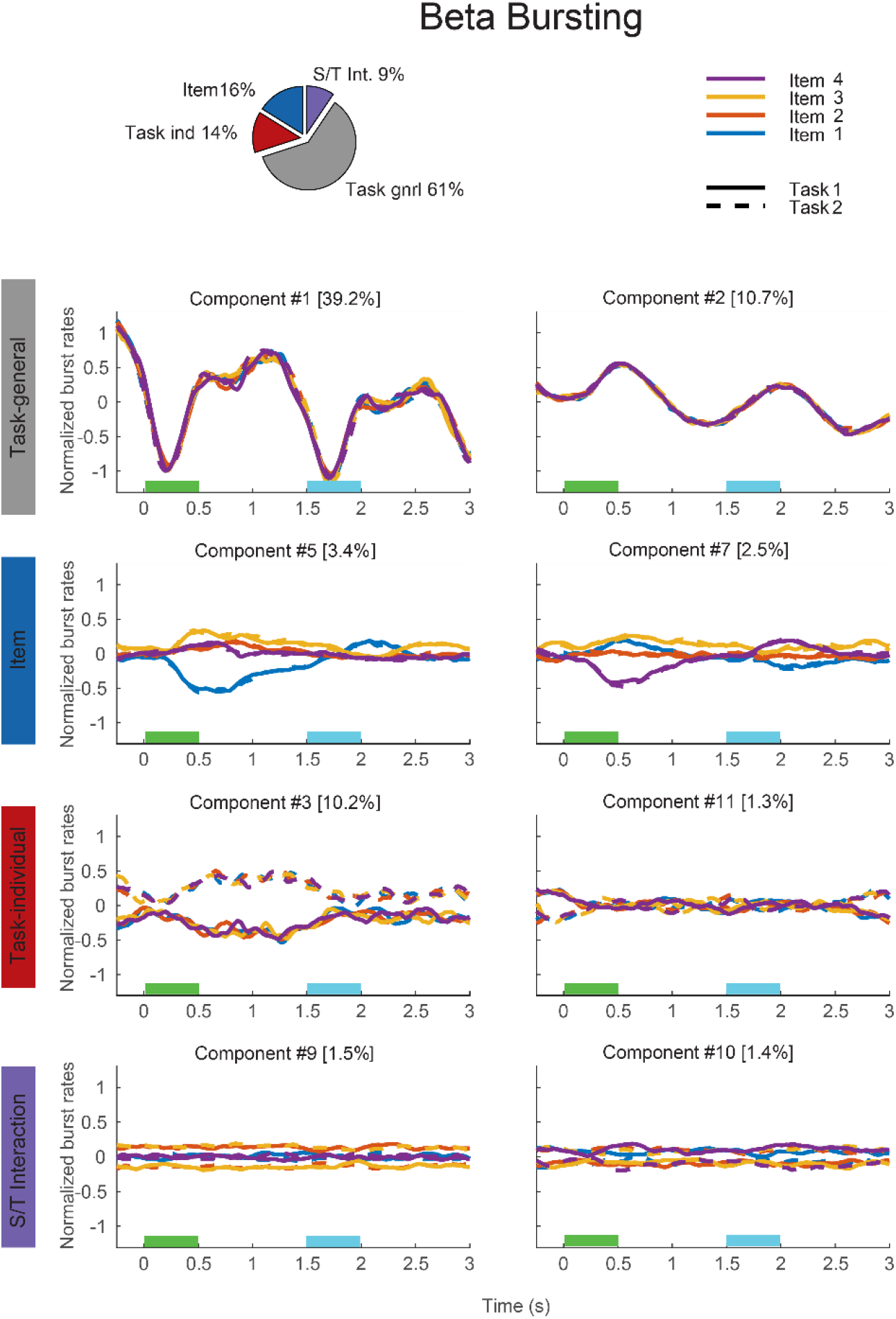
dPCA analysis of beta bursting. Same as in Figure 3 but for calculated over patterns of beta bursting.

**Figure S5.**
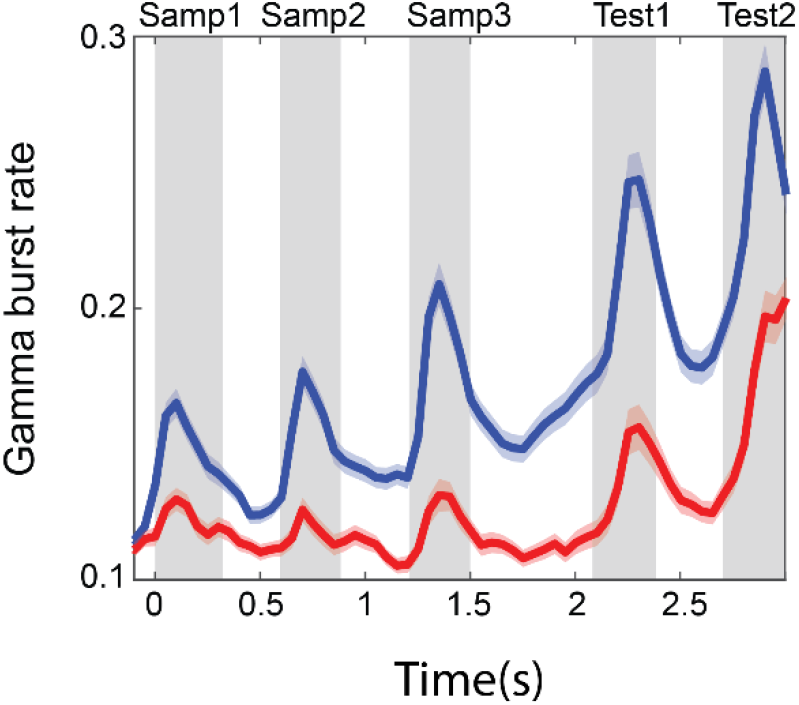
Same as Figure 4B, but for gamma bursting.

## Acknowledgements

We would like to thank the ERC starting grant 949131, the Swedish Research Council Starting Grant 2018-04197 and 2018-05360, the 2017 Young Investigator Grant from the Brain & Behavior Research Foundation, the National Institutes of Mental Health Grant R37MH087027 and 5K99MH116100-02, Office of Naval Research Multidisciplinary University Research Initiatives Grant N00014-16-1-2832, Swedish e-science Research Center for their support.

